# Iron deposition and functional connectivity alterations in the right substantia nigra of adult males with autism

**DOI:** 10.1101/2025.04.18.649491

**Authors:** Takashi Itahashi, Kazuyo Tanji, Yumi Shikauchi, Taiga Naoe, Tsukasa Okimura, Motoaki Nakamura, Haruhisa Ohta, Ryu-ichiro Hashimoto

## Abstract

The substantia nigra (SN) is a midbrain nucleus implicated not only in motor control and reward processing but also in higher-order cognitive functions. Iron homeostasis in this region is essential for neurotransmitter synthesis, especially for dopamine, and thus, iron dysregulation may contribute to the symptomatology of autism spectrum disorder (ASD). However, iron deposition and functional circuits of the SN in the autistic brain remain underexplored. This study investigated iron deposition and functional connectivity (FC) of the SN in 53 adult males with ASD and 99 typically developing controls using quantitative susceptibility mapping and resting-state fMRI. Compared to controls, the ASD group exhibited higher magnetic susceptibility in the right SN, suggesting elevated iron deposition. Within the ASD group, higher iron deposition was associated with more severe socio-communicative deficits and reduced sensory-seeking behavior. Seed-based FC analyses further revealed that the ASD group exhibited stronger FC between the right SN and bilateral visual cortices and reduced FC with the right superior frontal gyrus. These results highlight the critical role of SN in the autistic brain and indicate that altered iron homeostasis in the SN may contribute to disruptions in the dopaminergic system that underlie the core symptoms of ASD.

## Introduction

The substantia nigra (SN) is a midbrain nucleus traditionally recognized for its role in motor control and reward processing; however, recent findings underscore its critical involvement in higher-order cognitive functions (Kamiński et al. 2018; Batten et al. 2024). This nucleus contains distinct neuronal populations; dopaminergic neurons provide a feedback loop to the striatum, and GABAergic neurons relay basal ganglia output to the thalamus (Halliday 2004). In addition to connections with the striatum, the SN connects anatomically with widespread cortical brain regions, including prefrontal, somatomotor, and occipital cortices (Zhang et al. 2017). Through these mechanisms and anatomical organization, the SN helps regulate cortical activity (Gu et al. 2021). Therefore, disruptions in SN may disturb the balance of cortical networks critical for sensory and cognitive processes, contributing to the underlying pathophysiological mechanisms of neurodevelopmental disorders.

Aberrant SN function has been implicated in neurodevelopmental disorders, such as autism spectrum disorder (ASD). In particular, the midbrain dopamine hypothesis postulates that dysregulation of midbrain dopaminergic pathways, particularly the nigrostriatal and mesocorticolimbic circuits, may contribute to core ASD features, including socio-communicative deficits and restricted and repetitive behaviors (Pavăl 2017; Kosillo and Bateup 2021; Pavăl and Micluția 2021). Supporting this hypothesis, alterations in this region have been reported in ASD (Seif et al. 2021). For example, prior post-mortem studies have demonstrated that autistic individuals exhibit age-related alterations in the volume of the SN, with reduced cell volume in autistic children and increased cell volume in autistic adults (Wegiel et al. 2014, 2015). These microscopic structural alterations in this region may be driven by multiple factors, including metabolic disruptions (e.g., iron dysregulation).

Iron is vital for normal brain development, supporting oxidative metabolism, myelination, and neurotransmitter synthesis (Hare et al. 2013; Bogdan et al. 2016). In subcortical structures, both iron overload and iron deficiency have been associated with cognitive impairments (Larsen et al. 2020; Spence et al. 2020). Aberrant iron homeostasis in the SN, thus, may contribute to the formation of the ASD symptomatology. While elevated iron accumulation in the SN is a well-known feature of neurodegenerative disorders such as Parkinson’s disease (Rouault 2013; Ward et al. 2014; Levi et al. 2024), research on altered brain iron deposition in ASD is still in its early stages. Consequently, characterizing iron deposition using advanced neuroimaging techniques is critical for understanding SN dysfunction in ASD.

Quantitative susceptibility mapping (QSM) is an advanced MRI technique that quantifies the magnitude and spatial distribution of magnetic susceptibility. The positive and negative magnetic susceptibility values reflect the magnetic susceptibility of paramagnetic substances (e.g., ferritin and deoxyhemoglobin) and diamagnetic substances (e.g., myelin and calcification), respectively (Haacke et al. 2015; Harada et al. 2022). Recent QSM studies have reported altered iron deposition in ASD. For instance, autistic children have significantly lower magnetic susceptibility (i.e., reduced iron deposition) in the SN than typically developing controls (Tang et al. 2021, 2022; Zhou et al. 2025). Such iron deficiency in the SN could disrupt dopaminergic neurotransmission synthesis and neural development, potentially exacerbating socio-communicative deficits observed in ASD (Pavăl 2017). However, whether the reduced SN iron deposition observed in autistic children persists into adulthood or undergoes developmental changes remains unclear.

The current study aimed to examine both the structural and functional involvement of the SN in adults with ASD. We used QSM to quantify iron deposition in the SN of adults with ASD and examined its association with clinical symptoms. In addition, we analyzed resting-state functional connectivity (FC) of the SN with cortical regions to determine whether adults with ASD exhibit alterations in the SN-related functional circuitry. This multi-modal approach aims to investigate the association between SN abnormalities and ASD symptomatology. Our findings could provide new insight into midbrain contributions to ASD and have the potential to refine theoretical models of how subcortical systems shape the core features of autism.

## Materials and Methods

### Ethics statement

All participants provided written informed consent. The institutional review board at Showa University Karasuyama Hospital approved recruitment procedures and experimental protocols, which were conducted in accordance with the Declaration of Helsinki.

### Participants

In this study, we obtained MRI data and self-report questionnaires from 59 adult males with ASD and 103 typically developing control adult males (TDCs). All individuals with ASD were recruited from the outpatient unit of Showa University Karasuyama Hospital between July 2019 and March 2024, while TDCs were recruited via an advertisement that included information such as a brief summary of the research objectives, inclusion criteria (e.g., age ≧ 18 years old) and exclusion criteria (e.g., a history of psychiatric or neurological problems and no relatives who received the formal diagnosis of psychiatric disorders). Similar to our previous studies (Hashimoto et al. 2024; Itahashi et al. 2024), our clinical team assessed autistic participants’ developmental history, present illness, life history, and family history. The clinical team then made clinical diagnoses based on the DSM-5 (American Psychiatric Association 2013). Due to either poor image reconstruction, brain anomalies (e.g., cyst and cavernous hemangioma), or the incompleted questionnaires (e.g., Autism-Spectrum Quotient [AQ]), we excluded 6 participants with ASD and 4 TDCs from subsequent analyses.

Trained psychologists administered the second edition of the Autism Diagnostic Observation Schedule (ADOS-2) Module-4 (Gotham et al. 2007, 2009) to participants with ASD, and we confirmed that all the participants with ASD satisfied the cut-off scores (autism: *n* = 17 and ASD: *n* = 36). To estimate the full-scale intelligence quotient (IQ) score for each participant with ASD, the Wechsler Adult Intelligence Scale Third Edition or Fourth Edition was administered. On the other hand, the IQ scores of TDCs were estimated using the Japanese version of the National Reading Test (Matsuoka et al. 2006). All the participants completed the Japanese version of AQ (Baron-Cohen et al. 2001), Social Responsiveness Scale Second Edition (SRS-2) (Constantino 2012), Adolescent/Adult Sensory Profile (AASP) (Brown and Dunn 2002), and Edinburgh Handedness Inventory (Oldfield 1971). Since the Japanese version of SRS-2 is still under development and does not yet provide *T*-scores for total and subscale scores, this study only dealt with raw total scores. In the ASD group, 26 out of 53 participants with ASD were using either one or more of the following medications: antidepressants (*n* = 6), anti-anxiety drugs (*n* =3), sleeping pills (*n* =6), antipsychotic drugs (*n* = 5), stimulants (*n* =7), and antiepileptic drug (*n* = 2). We summarized the demographic and clinical information in Table 1.

**Table 1.** Summary of the demographic and clinical information.

|  | ASD | TDC |  |
| --- | --- | --- | --- |
|  | Mean (SD) | Mean (SD) | Group difference |
| Age | 27.57 (7.08) | 26.63 (6.30) | $t(150) = 1.29, p = 0.20$ |
| Estimated IQ | 108.13 (7.28) | 109.10 (7.76) | $t(150) = -0.75, p = 0.45$ |
| FIQ | 108.17 (13.99) | - |  |
| VIQ | 112.72 (14.30) | - |  |
| PIQ | 104.91 (16.00) | - |  |
| Handedness | 76.40 (47.30) | 69.35 (55.99) | $t(150) = 0.78, p = 0.44$ |
| AQ Total score | 32.68 (6.74) | 19.45 (7.21) | $t(150) = 11.02, p < 0.001$ |
| SRS-2 Total score | 107.53 (33.11) | 58.23 (26.33) | $t(150) = 10.03, p < 0.001$ |
| AASP |  |  |  |
| Low registration | 38.72 (11.40) | 26.53 (7.89) | $t(150) = 7.74, p < 0.001$ |
| Sensory seeking | 34.64 (9.57) | 36.05 (8.71) | $t(150) = -0.92, p = 0.36$ |
| Sensory sensitivity | 40.45 (11.88) | 32.10 (9.21) | $t(150) = 4.80, p < 0.001$ |
| Sensation avoidance | 42.28 (9.60) | 32.90 (8.64) | $t(150) = 6.13, p < 0.001$ |
| ADOS |  |  |  |
| Communication | 3.04 (1.00) | - | - |
| Reciprocal Social<br>Interaction | 6.02 (1.25) | - | - |
| Total score | 9.06 (1.95) | - | - |

Abbreviations: AASP: Adolescent/Adult Sensory Profile, ADOS: Autism Diagnostic Observation Schedule, AQ: Autism-Spectrum Quotient, IQ: intellectual quotient, SD: standard deviation, and SRS-2: social responsiveness scale, second edition

### MRI data acquisition

All the MRI data were acquired using a 3.0 Tesla MRI scanner (MAGNETOM Skyra fit; Siemens Healthcare, Erlangen, Germany) with a 32-ch head coil. The details of the scanning protocol were described elsewhere (Koike et al. 2021). The T1-weighted image was acquired with MPRAGE sequence (repetition time [TR]: 2500 ms, echo time [TE]: 2.18 ms, Inversion time [TI]: 1000ms, field of view [FoV]: 256 × 240 × 179.2 mm, resolution: 0.8 × 0.8 × 0.8 mm, slice oversampling: 7.1 %, flip angle: 8 degrees, partial Fourier: 6/8, GRAPPA factor: 2, Prescan normalize: ON). The T2-weighted image was acquired with SPACE sequence (TR: 3200 ms, TE: 564 ms, FoV: 256 × 240 × 179.2 mm, resolution: 0.8 × 0.8 × 0.8 mm, slice oversampling: 7.1%, GRAPPA factor: 2, Prescan normalize: ON). QSM image was acquired with a three-dimensional multi-echo gradient echo sequence (TR: 44 ms, TE1: 3.6 ms, TE2: 9.51 ms, TE3: 15.43 ms, TE4: 21.34 ms, TE5: 27.26 ms, TE6: 33.17 ms, TE7: 39.09 ms, FoV:240 × 240 mm, matrix size: 256 × 256, in-plane resolution: 0.9 × 0.9 mm, slice thickness: 1 mm, flip angle: 15 degrees, Phase partial Fourier: 6/8, GRAPPA factor: 2, Prescan normalize: ON). Resting-state fMRI (R-fMRI) data were acquired with a multi-band echo planar imaging sequence (TR: 800 ms, TE: 34.4 ms, flip angle: 52 degrees, FoV: 206 × 206 mm, matrix size: 86 × 86, in-plane resolution: 2.4 × 2.4 mm, slice thickness: 2.4 mm without gap, number of slices: 60, multi-band factor: 6, Prescan normalize: ON). We acquired two 5-minute R-fMRI for each phase encoding direction (i.e., anterior-to-posterior and posterior-to-anterior), yielding 20-minute R-fMRI data for each participant. During the R-fMRI runs, participants were asked to fixate their eyes on a cross-hair. We also acquired a spin-echo map for each phase encoding direction (TR: 6100 ms, TE: 60 ms, FoV: 206 × 206 × 144 mm, resolution: 2.4 × 2.4 × 2.4 mm, partial Fourier: OFF, Prescan normalize: ON) to enable distortion correction (Andersson et al. 2003).

### Structural MRI data preprocessing

The structural MRI and resting-state fMRI data were preprocessed using the Human Connectome Project (HCP) pipeline (Glasser et al. 2013) v4.3.0 (https://github.com/Washington-University/HCPpipelines). The details of the preprocessing steps were described elsewhere (Glasser et al. 2013). Briefly, anatomical images (i.e., T1-weighted and T2-weighted images) were fed into non-linear registration to the Montreal Neurological Institute (MNI) space and used for cortical surface reconstruction using FreeSurfer v6.3.0 (Fischl 2012) surface registration using multi-modal surface matching (Robinson et al. 2018). This procedure was followed by creating a myelin map, which was subsequently used for MSMAll registration, using T1-weighted divided by T2-weighted images and surface mapping (Glasser and Van Essen 2011). We used the Brain Connectomics Imaging Library (https://github.com/RIKEN-BCIL/bcil) to visually inspect the quality of anatomical images.

### QSM preprocessing

QSM images were reconstructed using QSMxT (Stewart et al. 2022) v7.2.0 (https://qsmxt.github.io/QSMxT/), available as a container via NeuroDesk (Renton et al. 2024). The details of the preprocessing steps are described elsewhere (Stewart et al. 2022). Briefly, QSMxT integrated and automated phase unwarping using a rapid open-source minimum spanning tree algorithm (Dymerska et al. 2021), background field removal with projection onto dipole fields (Liu et al. 2011), and sparsity-based rapid two-step dipole inversion (Kames et al. 2018). This pipeline was congruent with recent consensus statement recommendations for best-practice QSM reconstruction (QSM Consensus Organization Committee et al. 2024). QSMxt enabled a novel two-pass combination method for hole-filling and artifact reduction, which performed parallel QSM masking and reconstruction on susceptibility sources identified as reliable and less reliable. Dual QSM images were combined into a final integrated image that was more robust to reconstruction errors and streaking artifacts than those produced using a single-pass approach. After reconstruction, we visually inspected all constructed susceptibility maps and excluded if there were any apparent artifacts.

We further mapped the obtained quantitative susceptibility map (i.e., Chimap) onto the fsLR32k space using the FMRIB Software Library (FSL) (Smith et al. 2004; Jenkinson et al. 2012) and Connectome workbench (https://github.com/Washington-University/workbench).

The magnetic susceptibility map was registered to the structural T1-weighted AC-PC space using the first echo magnitude image and the white surface using FSL’s FLIRT (Jenkinson and Smith 2001; Jenkinson et al. 2002) and FreeSurfer’s bbregister (Greve and Fischl 2009) and then mapped onto the cortical surface using an algorithm weighted towards the cortical mid-thickness (Glasser and Van Essen 2011). For each mid-thickness surface vertex on the native mesh, the algorithm identified cortical ribbon voxels within a cylinder orthogonal to the local surface. The surface maps were resampled based on MSMAll surface registration (Robinson et al. 2014; Glasser et al. 2016) and onto the 32k group average surface mesh. For statistical analyses, we extracted mean magnetic susceptibility values of the bilateral SN using the brainstem navigator (Bianciardi et al. 2015).

### R-fMRI data preprocessing and seed-based FC analyses

R-fMRI data was also preprocessed using HCP pipeline v4.3.0 (Glasser et al. 2013). R-fMRI data were first corrected for distortion and head motion. The distortion from B0 static field inhomogeneity was corrected using opposite phase encoding spin echo fieldmap data using TOPUP (Andersson et al. 2003). It was then warped and resampled to MNI space at a 2 mm resolution. The region of the cortical ribbon in the fMRI volume was further mapped onto the cortical surface and combined with voxels in the subcortical gray region to create 32k grayordinates. Four runs of the R-fMRI data were merged and fed into multi-run ICA-FIX (Salimi-Khorshidi et al. 2014). After automated noise classification, all independent components underwent visual inspection and were manually reclassified. The denoised R-fMRI data, in combination with myelin and cortical thickness, was further used for multi-modal registrations over the cortical surface, followed by removing registration bias after multimodal registration (Glasser et al. 2016).

To remove the effects of nuisance signals from white matter (WM) and cerebrospinal fluid (CSF) that still remained, we performed additional post-processing in the following manner. We first used aComCor to extract artifactual signals from the union of WM and CSF masks (Behzadi et al. 2007). In this study, the aCompCor method extracted five principal components as artifactual signals. In addition to these signals, we used 24 head motion parameters and a global signal as nuisance regressors. After regressing the effects of these nuisance variables, a band-pass filter (0.01–0.08 Hz) was applied. We calculated frame-wise displacement (FD) and excluded time points from subsequent analyses if their FD values exceeded 0.5 mm (Power et al. 2012, 2014).

We further investigated whether the ASD group exhibited alterations in the functional pathways seeding from the SN to cortical regions. We used the HCP-MMP v1.0 cortical atlas (Glasser et al. 2016) as ROIs. Pearson’s correlation coefficients between the right SN and other ROIs were computed, and Fisher’s *r*-to-*z* transform was applied, yielding a 360-dimensional connectivity vector for each participant.

We computed the mean FD for each participant and then identified participants who showed excessive head motions during the scans using the interquartile range method. Using this outlier method, the threshold for the mean FD was 0.21 mm, and we excluded 6 participants (ASD: *n* = 4 and TDC: *n* = 2) from the subsequent analyses (i.e., ASD: *n* = 49 and TDC: *n* = 97).

## Statistical analysis

### Group comparisons of magnetic susceptibility values in the SN

We used a general linear model, with age and medication status as covariates of no interest, to test whether the ASD group exhibited altered magnetic susceptibility values compared to the TDC group. We constructed a general linear model with the function “*fitlme*” implemented in MATLAB R2021b (Mathworks). The statistical threshold was set to *p* < 0.05 after correcting the family-wise error rate (FWER) using the Bonferroni correction.

Once statistically significant between-group differences were observed, we examined the associations between altered magnetic susceptibility values and the severity of clinical symptoms assessed by the self-report questionnaires (i.e., AQ, AASP, and SRS-2) and ADOS using partial correlation analyses. Age and medication status were included as covariates of no interest. The statistical threshold was set to *p* < 0.05 after false discovery rate (FDR) correction (Storey 2002).

### Group comparisons of the SN-related functional pathways

To examine whether the ASD group exhibited altered FCs compared to the TDC group, we used a general linear model with age, medication status, and mean FD as covariates of no interest. The statistical threshold was set to *p* < 0.05 after FDR correction (Storey 2002).

To confirm whether altered FCs were associated with the severity of clinical symptoms within the ASD group, we further performed partial correlation analyses, controlling for age, medication status, and mean FD. The statistical threshold was set to *p* < 0.05 after FDR correction (Storey 2002).

## Results

### Altered magnetic susceptibility in the right SN

As shown in Figure 1, statistical analyses revealed that the ASD group exhibited higher magnetic susceptibility values in the right SN after controlling for age and medication (*t*-value = 2.61, *p*_FWER_ = 0.020). At the same time, those in the left SN could not survive after FWER correction (*t*-value = 2.21, *p*_FWER_ = 0.058). For the nuisance covariates, the effect of age was statistically significant in the bilateral SN (left SN: *t*-value = 2.78, *p* = 0.006; right SN: *t*-value = 3.36, *p* < 0.001); the effect of medication was also statistically significant in the right SN (*t*-value = −2.01, *p* = 0.047), but not in the left SN (*t*-value = 0.34, *p* = 0.734).

**Figure 1.**
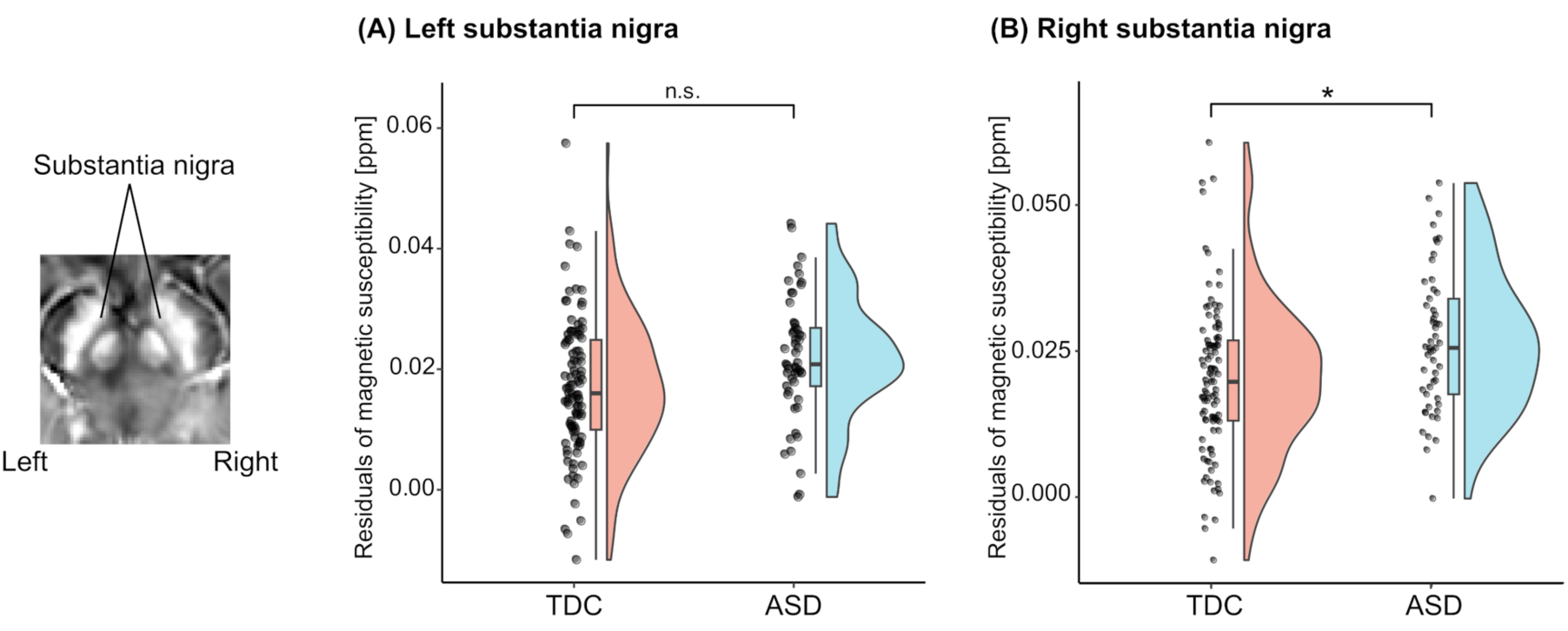
Group comparisons of magnetic susceptibility in the bilateral SN. (A) Raincloud plots of magnetic susceptibility in the left SN. (B) Raincloud plots of magnetic susceptibility in the right SN. The asterisk indicates statistical significance after correcting the family-wise error rate (FWER; i.e., *p_FWER_* < 0.05), while the term “n.s.” indicates *p_FWER_* > 0.05.

We repeated statistical analyses using undersampling procedures to rule out the possibility that our findings were due to the imbalance between the two groups. We randomly selected 53 TDCs and then examined the effect of diagnosis while controlling for age and medication status. We counted the number of times *p*-values were below 0.05 after FWER correction. We repeated this procedure 1,000 times and found that the statistically significant effect of diagnosis was observed 658 times (803 times without FWER correction) in the right SN. Such an effect was observed 494 times (549 times without FWER correction) in the left SN. Binomial tests further confirmed that the occurrence of a statistically significant effect of diagnosis in the right SN was greater than chance (*p* < 0.001). These results support that our findings were robust to the imbalanced number of participants between the two groups.

### Associations between altered magnetic susceptibility in the right SN and the clinical symptoms

Partial correlation analyses revealed that, within the ASD group, atypical magnetic susceptibility values in the right SN statistically significantly correlated with the ADOS “Communication” subscale scores (*ρ* = 0.37, *p*_FDR_ = 0.037), the ADOS total scores (*ρ* = 0.36, *p*_FDR_ = 0.037), and AASP sensory seeking scores (*ρ* = −0.40, *p*_FDR_ = 0.037) (see Figure 2). However, other scores (i.e., AQ total score, AASP low registration, AASP sensory sensitivity, AASP sensation avoidance, and SRS-2 total score) did not reach the statistical threshold (all *p*_FDR_ > 0.69).

**Figure 2.**
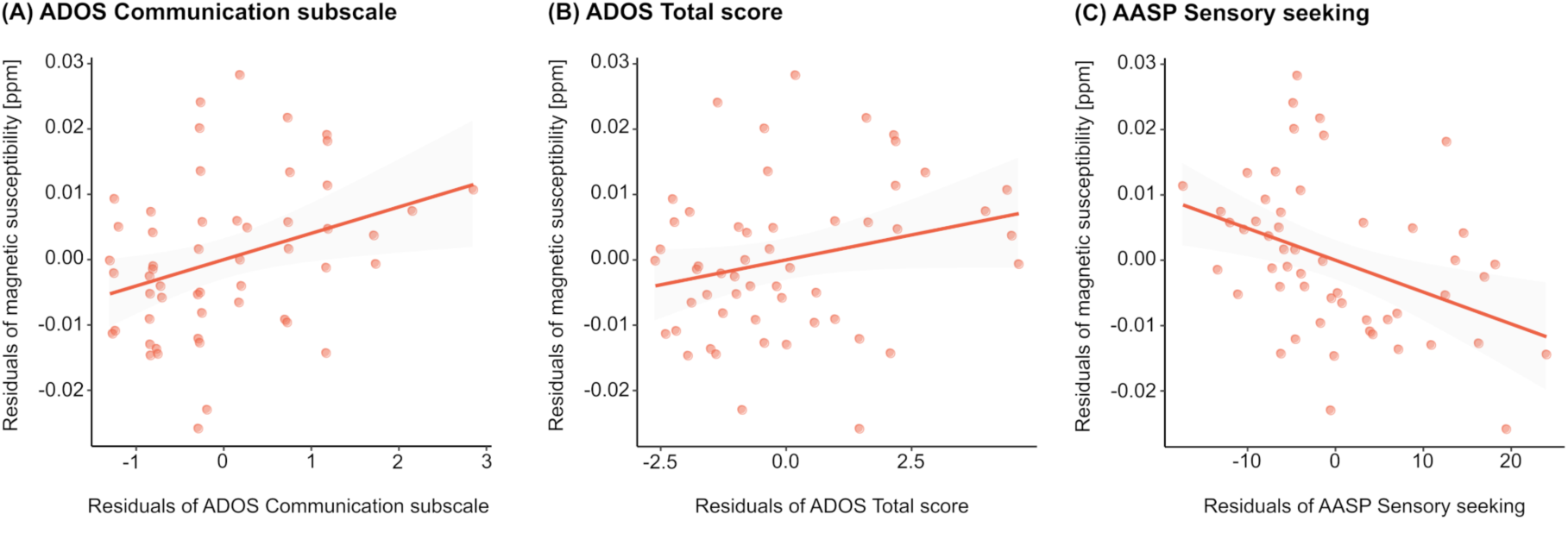
Associations with the severity of autistic symptoms and the magnetic susceptibility value of the right SN within the autistic group. (A) ADOS Communication subscale. (B) ADOS Total score. (C) AASP Sensory seeking score. The effects of age and medication status were removed from both the magnetic susceptibility value and the severity of autistic symptoms.

### Altered right SN-related functional connectivity

As shown in Figure 3, compared with the TDC group, the ASD group exhibited stronger FCs between the right SN and the bilateral visual cortices, including the visual area 1 (V1), V2, V3A, V6, and the ventromedial visual area 1 (VMV1). In addition, the ASD group showed a weaker FC between the right SN and the right superior frontal gyrus (SFG; p10p) (*p*_FDR_<0.05).

**Figure 3.**
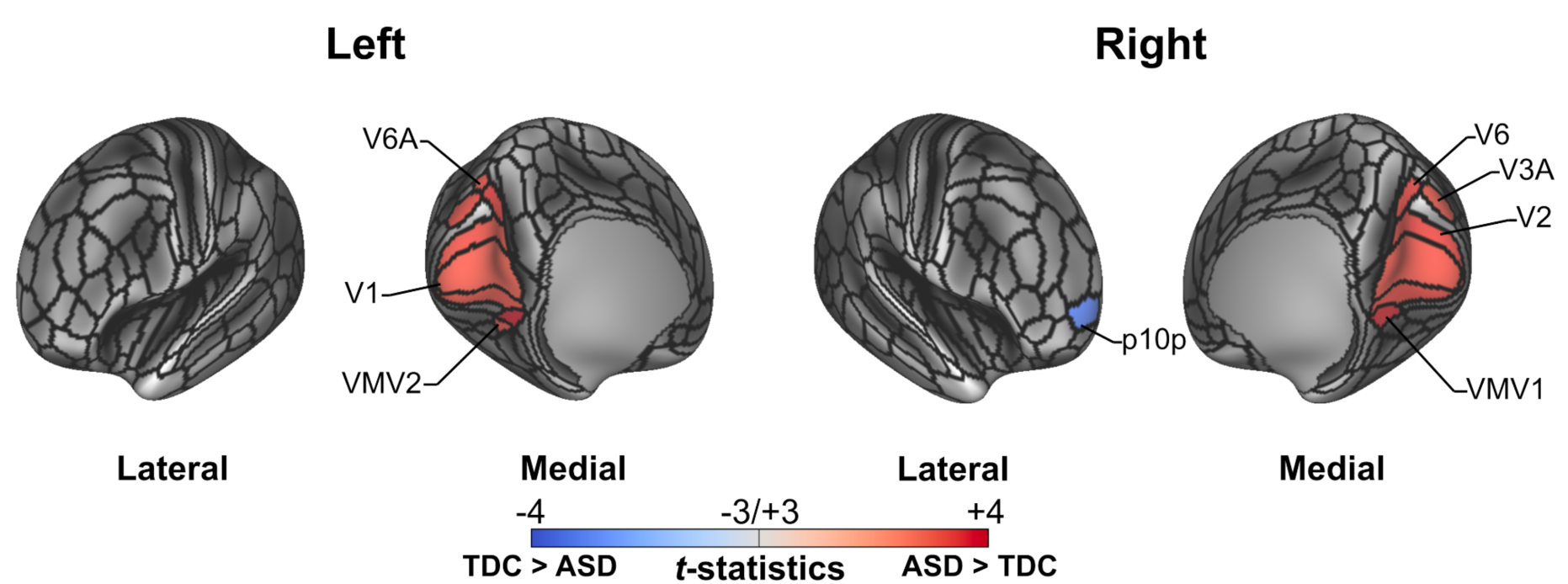
Comparisons of the right SN-related FCs between the two groups. The red and blue colors show stronger and weaker FCs in the ASD group. The black-colored boundaries represent the HCP-MMP v1.0 cortical parcels. The names of each parcel follow the HCP-MMP v1.0 notation.

Within the ASD group, no associations between altered FCs and the clinical symptoms survive after correcting multiple comparisons (all *p_FDR_* > 0.73). With a liberal statistical threshold (i.e., *p* < 0.05), we observed that the strengths of FCs between right SN and left V1 and left VMV1 were positively correlated with the AASP sensory seeking scores (left V1: *ρ* = 0.314, *p* = 0.034, and left VMV1: *ρ* = 0.381, *p* = 0.009), while those between the right SN and left VMV1 and left VMV2 were negatively correlated with AASP avoidance scores (left VMV1: *ρ* = −0.308, *p* = 0.037, and left VMV2: *ρ* = −0.375, *p* = 0.010).

## Discussion

This study investigated the involvement of SN in adult males with ASD using QSM and R-fMRI. Compared to the TDC group, the ASD group exhibited higher magnetic susceptibility values in the right SN, providing evidence for elevated brain iron deposition. Within the ASD group, the magnitude of magnetic susceptibility was significantly associated with the severity of socio-communicative deficits measured by ADOS-2 and sensory-seeking scores measured by AASP. Moreover, the ASD group exhibited stronger FCs with visual cortices and a weaker FC with the right SFG. These findings suggest that SN may play a critical role in the pathophysiological mechanisms of ASD.

The elevated magnetic susceptibility in the right SN of adults with ASD indicates accelerated iron deposition, which may disrupt dopaminergic neurotransmitter synthesis. Statistical analyses controlling for age and medication status have confirmed that the effect of diagnosis is robust, as undersampling procedures consistently demonstrated a significant diagnostic difference in the right SN. In contrast to our findings in adults with ASD, prior QSM studies have shown reduced magnetic susceptibility in autistic children (Tang et al. 2021, 2022; Zhou et al. 2025). Although the precise reasons for the opposite iron deposition pattern observed in autistic children and adults remain unclear, several developmental factors might contribute to this phenomenon. The dopaminergic system and brain iron regulation undergo significant maturation and reorganization throughout development (Ernst et al. 2009; Larsen et al. 2020). Additionally, autistic individuals exhibit atypical neurodevelopmental trajectories characterized by altered cortical maturation and connectivity patterns (Wallace et al. 2015; Andrews et al. 2021; Lee et al. 2025). Thus, the complex interplay between the maturational process and iron metabolism might explain why opposite patterns of iron deposition emerge in autistic children compared to autistic adults. Future longitudinal studies are necessary to better characterize developmental trajectories of iron deposition and clarify their functional implications in ASD across the lifespan.

In addition to elevated iron deposition, the higher magnetic susceptibility values in the right SN were significantly correlated with more severe socio-communicative deficits in autistic adults. Iron overload could induce dopaminergic dysfunction through oxidative stress-mediated mechanisms, as observed in neurodegenerative disorders (Rouault 2013). Furthermore, dysfunctions in midbrain dopaminergic pathways, particularly the nigrostriatal and mesocorticolimbic circuits, have been implicated in core ASD features (Pavăl 2017; Pavăl and Micluția 2021). Although our study did not directly assess dopamine levels, the association between increased iron deposition in the SN and greater socio-communicative deficits is consistent with the hypothesis that dopaminergic dysregulation underlies these clinical manifestations (Kohls et al. 2012). Furthermore, we observed a significant correlation between elevated magnetic susceptibility in the right SN and reduced sensory-seeking behavior in autistic adults. Given previous evidence linking atypical sensory seeking to impaired adaptive functioning (Baranek et al. 2018; Worthley et al. 2024; Itahashi et al. 2025), further investigation is needed to investigate a potential triad association between brain iron deposition, sensory seeking, and executive functions in autism.

Our seed-based FC analysis demonstrated that the ASD group exhibited increased FCs between the right SN and visual cortices and a reduced FC to the right SFG. Given that the SN anatomically connects with widespread cortical regions, including prefrontal, somatomotor, and occipital cortices (Zhang et al. 2017), and plays a key role in orchestrating cortical activity (Gu et al. 2021), our findings may be associated with prior reports of an atypical functional connectome hierarchy in ASD. In particular, recent studies have documented an acceleration of stepwise connectivity (Hong et al. 2019) and a shorter intrinsic neural timescale in the visual cortex (Watanabe et al. 2019), both of which may reflect a shift in the balance between bottom-up sensory processing and top-down cognitive regulation. Although correlations between atypical FCs and sensory symptoms did not survive multiple comparison corrections, there was a trend toward associations with sensory seeking and sensory avoidance. Taken together, our findings in atypical SN-related FCs likely contribute to the atypical sensory experiences observed in ASD. Future research should explore how these SN-related connectivity abnormalities impact the information-processing hierarchy underlying sensory and cognitive function integration in autism.

Several methodological considerations temper the interpretation of our results. First, our sample consisted exclusively of adult males with ASD, which limits the generalizability of our findings. ASD is heterogeneous owing to sex and developmental stages; females with ASD can differ in neurobiology and behavior (Lai et al. 2013; Trakoshis et al. 2020). Future studies should include female participants to determine whether the increased SN iron and connectivity are universal findings in adults or specific to adult males. Second, we did not consider the subregions of the SN. The SN consists of SN pars compacta (SNc) and SN pars reticulata (SNr), which contain different neural populations—dopaminergic and GABAergic neurons, respectively (Halliday 2004). These subregions have distinct anatomical and functional roles: the SNc is primarily involved in modulating striatal activity through dopaminergic projections, while the SNr serves as a major output nucleus of the basal ganglia via GABAergic transmission (Halliday 2004). By not differentiating between these subregions, our analysis may have overlooked region-specific alterations in iron deposition and connectivity that could provide deeper insights into the neurobiology of ASD. Future studies should evaluate the SNc and SNr, clarifying their contributions to ASD pathophysiology. Third, although QSM is an advanced technique for assessing tissue iron content, it does not directly measure dopaminergic function. As a result, our study cannot conclusively link elevated iron deposition with dysregulated dopamine signaling. Future studies should consider other imaging modalities, such as neuromelanin-sensitive MRI (Cassidy et al. 2019; Kuai et al. 2024) or positron emission tomography of dopamine receptors (Kubota et al. 2020; Murayama et al. 2022), to assess the relationship between iron homeostasis and dopaminergic activity in ASD.

In summary, this study demonstrates that adult males with ASD exhibit both structural and functional abnormalities in the SN. Elevated iron deposition in the right SN and its association with socio-communicative deficits and sensory-seeking behaviors support the hypothesis that dopaminergic dysregulation contributes to ASD pathophysiology. Moreover, altered FC patterns, characterized by increased FCs with visual cortices and reduced FC with the right SFG, underscore disruptions in the balance between sensory processing and executive control. These results highlight the critical role of the SN and emphasize the need for further investigation into midbrain circuit dysfunction in autism.

## Acknowledgments

We would like to express our gratitude to Noriko Ishimura, Mika Kato, and Taku Sato for their assistance in recruiting participants for this study.

## CRediT statement

Takashi Itahashi (Conceptualization, Investigation, Data curation, Formal analysis, Funding acquisition, Methodology, Visualization, Writing – original draft, Writing – review & editing), Kazuyo Tanji (Funding acquisition, Writing – review & editing), Yumi Shikauchi (Writing – review & editing), Taiga Naoe (Writing – review & editing), Tsukasa Okimura (Investigation, Writing – review & editing), Motoaki Nakamura (Investigation, Funding acquisition, Writing – review & editing), Haruhisa Ohta (Investigation, Writing – review & editing), Ryu-ichiro Hashimoto (Conceptualization, Funding acquisition, Writing – review & editing). All authors have read and approved the manuscript.

## Declaration

The authors declare no competing interests.

